# Computationally reconstructing the evolution of cancer progression risk

**DOI:** 10.1101/2024.12.23.629914

**Authors:** Kefan Cao, Russell Schwartz

**Author notes:** Address for correspondence. Russell Schwartz, Carnegie Mellon University, Pittsburgh, PA 15213 USA.

## Abstract

Understanding the evolution of cancer in its early stages is critical to identifying key drivers of cancer progression and developing better early diagnostics or prophylactic treatments. Early cancer is difficult to observe, though, since it is generally asymptomatic until extensive genetic damage has accumulated. In this study, we develop a computational approach to infer how once-healthy cells enter into and become committed to a pathway of aggressive cancer. We accomplish this through a strategy of using tumor phylogenetics to look backwards in time to earlier stages of tumor development combined with machine learning to infer how progression risk changes over those stages. We apply this paradigm to point mutation data from a set of cohorts from the Cancer Genome Atlas (TCGA) to formulate models of how progression risk evolves from the earliest stages of tumor growth, as well as how this evolution varies within and between cohorts. The results suggest general mechanisms by which risk develops as a cell population commits to aggressive cancer, but with significant variability between cohorts and individuals. These results imply limits to the potential for earlier diagnosis and intervention while also providing grounds for hope in extending these beyond current practice.

**Availability:** The code used to conduct the analysis is available at: https://github.com/kefanc2/CancerRisk

## 1. Introduction

Cancer remains one of the major sources of mortality globally (Frick et al., 2023; Zhang et al., 2023) despite many years of intensive research into prevention and treatment. Great hope for reductions in mortality has been placed in earlier screening, to identify more cancers when they are treatable (Frick et al., 2023). While such efforts have saved lives, they underperformed early hopes in part due to overtreatment (Bhatt and Klotz, 2016): the statistical conclusion from reductions in mortality that many cancers detected early would never have been threatening to their hosts. Concern about the harm to patients of overtreatment in turn has led to undertreatment (Kocik, 2020), as evolving clinical practice to take more cautious approaches to cancer treatment of cancers judged unlikely to be aggressive can lead to failure to aggressively treat some that need it. The challenge of navigating the complementary issues of overtreatment and undertreatment has led to numerous efforts to more accurately distinguish threatening from non-threatening cancers (Bassel et al., 2022; Savalia and Verma, 2023; Kartikasari et al., 2024), essentially a problem of better predicting future progression (Zhang et al., 2023).

Much insight into how tumors evolve and progress has come from the discipline of cancer phylogenetics (Burrell et al., 2013; Schwartz and Schäffer, 2017), i.e., the study of the evolutionary history of cancer cells. Phylogenetic analysis can reveal order of mutations in a cancer cell, the timing of these mutations, and the relationships between different subclones, e.g., whether the most deadly clones arise from a single lineage or through parallel events (Hong et al., 2015).

Work on cancer evolution has suggested that cancer is not normally a primary illness but rather usually a late stage of a process of somatic hypermutability, either intrinsic or due to environmental factors (Alexandrov et al., 2013), that could potentially be used to predict tumor risk long before the cancer becomes phenotypically abnormal Tao et al. (2021). Combined with the insight of overtreatment, that many such hypermutable lesions and even cancers will never actually become threatening to the patient, suggests that our primary goal in early cancer screening should not be to detect cancers *per se*, but to detect lesions of genetic damage that will go on to be threatening to the patient.

Solving such prediction problems has led to statistical inference and machine learning methods taking on a more prominent role in personalized and precision cancer treatment. See, for example, Masucci et al. (2024) for a recent review. This need to predict progression has led to major lines of inquiry into prediction from pathology imaging (Shaw et al., 2022; Unger and Kather, 2024), genomic data (Unger and Kather, 2024), and other clinical assays (Denninghoff, 2021). Such tools have yielded ever better ability to identify those cancers that will threaten patient lives and predict which tumors will likely respond to what treatments (Mechahougui et al., 2024). Nonetheless, they are limited by our ability to gather data about tumors, our imperfect understanding of the factors governing their biology, and by the inherent stochasticity of the progression process.

Our prospects for predicting future cancer progression hinge on the question of when a cell lineage becomes committed to being a cancer, or a cancer with a bad progression outcome. If risk of a cancer progressing is essentially constant until the moment it progresses, then the prediction task is essentially impossible. However, if risk of progression gradually increases over the history of a cancer then there is hope of predicting progression well before it occurs. There has been considerable prior work to answer variants of this question of how a tissue becomes committed to being an aggressive cancer. These include the classic two-hit model (Knudson Jr, 1971) and the more recent “bad luck” model (Tomasetti and Vogelstein, 2015) that suggests that a cancer’s fate is typically determined by a single chance mutation. These can be conceptualized as different ways of reasoning about how we expect measures of progression risk to vary over the history of a cell lineage on its way to any given progression outcome. For example, does risk of progression suddenly rise as from a single chance mutation shifting a tissue from low-risk to high-risk or does it increase gradually as from a series of mutations each slightly adding to aggressiveness? Do these changes tend to occur early in the tumor’s development, long before a cancer is likely detected, or late once it is already advanced? The answers to these questions may be of great practical importance in setting limits on the prospects for early detection of high-risk lesions or early interventions to keep incipient cancers off of high-risk pathways.

Our central goal in this study is to establish a model for reconstructing how progression risk develops over a tissue’s history as it transitions from healthy to cancerous and potentially to lethality. We want to ask, if we could have observed the cell populations that eventually become cancers from their earliest development, how early could we have identified that they were on a trajectory to aggressive cancer. We emphasize that our goal here is not to provide the definitive answer to this question, but to show 1) that it is theoretically possible to ask it by computational methods applied to feasible data sources and 2) that doing so will provide valuable insight into basic cancer research and potentially actionable knowledge for improving early cancer diagnosis and treatment. We recognize that we cannot definitively answer the question yet, largely because we do not have the ideal data for the proposed analysis, but proceed in the hope that this inspires future work to answer the question more definitively.

Our major contributions are:

- Developing a paradigm for applying tumor phylogenetics combined with machine learning to characterize how the landscape of cancer risk evolves over time.
- Implementing a realization of this paradigm using existing tools and data resources.
- Applying that realization to a pilot study of breast, lung, and colorectal cancers to suggest how cancer risk evolves and how it can vary between tumor types and individual patients.

The remainder of this paper describes how we accomplish these steps and examines the results and conclusions we can draw from them. Finally, in the Discussion, we return to considering how this question might be asked better in the future.

## 2. Method

At a high level, our approach is as follows. We begin by training a model to predict progression risk via Cox regression (Cox, 1972) on survival data from several TCGA cohorts (Cancer Genome Atlas Research Network et al., 2013). We then infer tumor phylogenies, using PhyloWGS (Deshwar et al., 2015), to infer a temporal model of progression of each tumor. We apply the risk model to inferred earlier states of progression to infer a profile of how risk evolved versus time on an individual basis. Finally, we analyze these models collectively to characterize overall population trends and their variability within and between cohorts. While we recognize that each stage of this process involves inferences that are prone to errors and uncertainty, we put it forth as a prototype for how to study the question of evolution of progression risk that might be improved upon in the future through better cohorts, data sources, and computational analyses. This general approach is summarized in Figure 1 and elaborated upon in the remainder of Section 2.

**Fig. 1:**
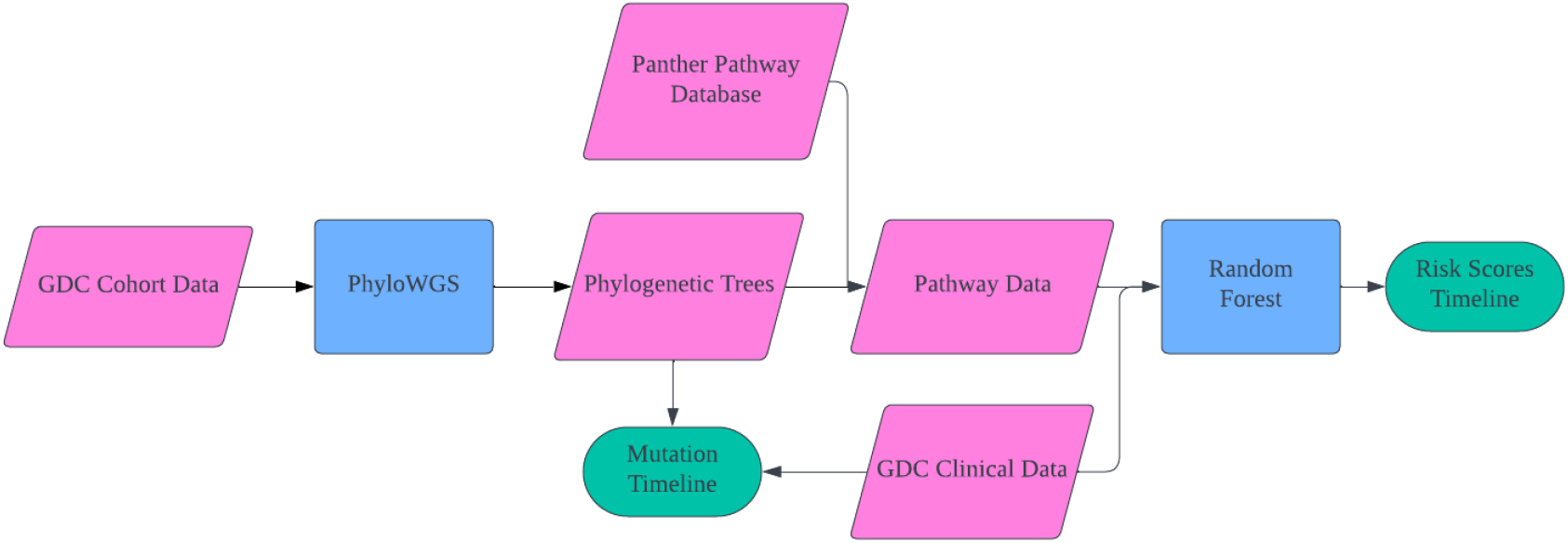
Summary figure describing the overall analysis pipeline.

### 2.1. Data Collection and Preprocessing

We apply our methodology to data from The Cancer Genome Atlas (TCGA) (Cancer Genome Atlas Research Network et al., 2013), which provides publicly available genomic data from large cohorts of thirty three different cancer types with associated clinical and demographic metadata. We accessed TCGA data through the Genomic Data Commons (GDC) data portal (Grossman et al., 2016), restricting analysis for the present study to DNA-seq single nucleotide variant (SNV) data. We also downloaded the clinical data for these cohorts to extract survival information of the patients. We restricted our analysis to cohorts with at least 400 subjects and low class imbalance in survival outcomes over the timeline of monitoring due to the need for predictive regression models of censored survival. We therefore chose to focus the study on two cancers: lung adenocarcinoma (TCGA-LUAD) and colorectal adenocarcinoma (TCGA-COAD).

### 2.2. Phylogenetic Analysis

A limitation of TCGA data for the present purpose is that it normally offers only one bulk DNA-seq sample per patient, which makes clonal phylogenetic inference challenging. To reconstruct the phylogenetic trees of the cohorts, we used PhyloWGS (Deshwar et al., 2015), a Bayesian method for tumor phylogeny inference from bulk SNV data, which our prior experience has shown to works comparatively well on single bulk samples compared to other popular methods. We applied PhyloWGS to the SNV data of each cohort to infer the evolution history of the cancer cell lineages. Where PhyloWGS inferred multiple possible trees, We selected the tree with the highest posterior probability as the final tree. We then used these trees to estimate the time points of key mutation in the evolution of cancer cells as described below. To ensure a reasonable runtime and exclude samples with insufficient mutations, we limited our analysis to samples with sizes ranging from 50KB to 2MB. Additionally, PhyloWGS failed on a small number of samples from both cohorts. Consequently, approximately 80% of the samples from each cohort were included in the study.

### 2.3. Pathway and Mutation Analysis

Since the number of mutations in the SNV data is large (≥ 10, 000 genes for 400 1100 patients), we took a systems approach to reduce the problem dimension by identifying key pathways relevant to tumor progression and aggregating mutations on each pathway Park et al. (2008). We used the Panther (Thomas et al., 2022; Mi and Thomas, 2009) database to convert raw SNV data to affected pathways, assuming that any SNV in a gene in a pathway affects the pathway. We then used the pathway data to identify the key pathways that are affected by the mutations. We also used random forest and Coxnet regression, via the scikit-survival (Pölsterl, 2020) package, applied to the pathway data to identify key pathways that affect patient survival.

## 3. Results

We also used random forest model to get risk scores for each patient based on the pathway information of the mutations. Since the set of variants and affected pathways for the two cancers are different, we trained two models for the two types of cancer respectively. Figure 2 shows how predicted risk varies over time for the two cohorts, LUAD (Fig 2a-b) and COAD (Fig 2c-d), separating subjects by survival outcome. Both cohorts show a qualitatively similar portrait of slowly increasing risk score over time. For both, mean risk scores are somewhat elevated in individuals with bad versus good survival outcomes throughout the tumor history, indicating that there are at least sometimes intrinsic differences predictive of outcome from the earliest stages of cancer development. However, the variability patient-to-patient is substantially larger than the difference between good and bad outcome subgroups. In the case of LUAD, there is minimal separation by outcome early on, but they begin to diverge in the latter half of their evolutionary trajectory, indicating that a significant portion of the determination of outcome occurs late in the tumor’s development. In the case of COAD, the separation between surviving and deceased subjects is more consistent across the tumor’s timeline.

**Fig. 2:**
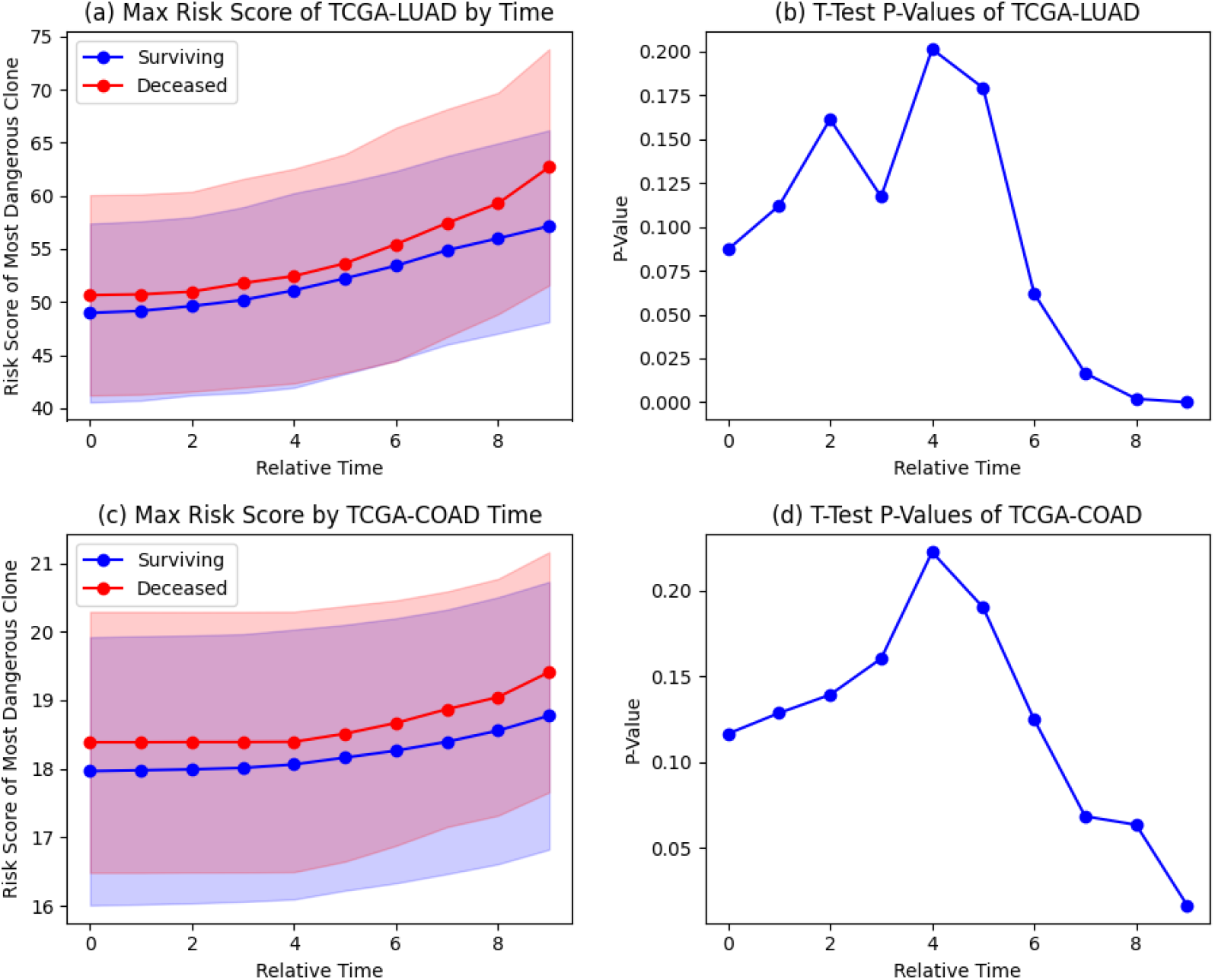
Risk scores and p-values versus inferred time for LUAD and COAD cohorts. (a) Risk score of the most high-risk clone vs. inferred time for LUAD subjects. (b) p-value for distinguishing by survival outcome in LUAD subjects. (c) Risk score of the most high-risk clone vs. inferred time for COAD subjects. (d) p-value for distinguishing by survival outcome in COAD subjects.

For both types of cancers, risk scores are significantly different between living and dead patients, as assessed by a t-test (p-value *<* 0.05). In both cases, however, ability to statistically distinguish patients by outcome improves sharply over the latter half of the trajectory as risk for both outcome groups increases. The separation only becomes statistically significant only fairly late in the progression process, although the time at which significance is achieved is in part a function of the cohort size among other study variables and not just an intrinsic property of the system.

To better understand what drives the evolution of the risk scores, we plotted for each tumor type the mean fraction of mutations of the five most frequently mutated driver genes for the highest-risk clones. These appear as Fig 3 (LUAD) and Fig 4 (COAD). In each figure, we provide two subplots to separate patients surviving over the course of the TCGA study followup from those deceased at the end of follow-up. To map mutations to an approximate time axis, we assumed a molecular clock hypothesis with mutations accumulating at a uniform rate across the tree and the length of a tree branch proportional to the number of mutations it contains. The figures show that while the overall risk evolution appears similar for the two cohorts, they exhibit quite different patterns of accumulation of key mutations in the evolution of cancer cells is different between the two types of cancers. The curves for LUAD involve a relatively more gradual increase in inferred mutations over the course of a tumor’s development. They are also less dominated by any one mutation, although TP53 emerges early as the most common mutation in each. There is little evident qualitative difference between the good and bad outcome plots aside from higher levels of TP53 mutation and lower levels of CSMD3 in bad-outcome cancers. COAD, by contrast, shows a sharper increase and higher rate of mutation, especially for the two most mutated genes (TP53 and APC). While mutations are predicted to accumulate in these genes from relatively early times, there is a notable increase in their mutation frequencies relatively late.

**Fig. 3:**
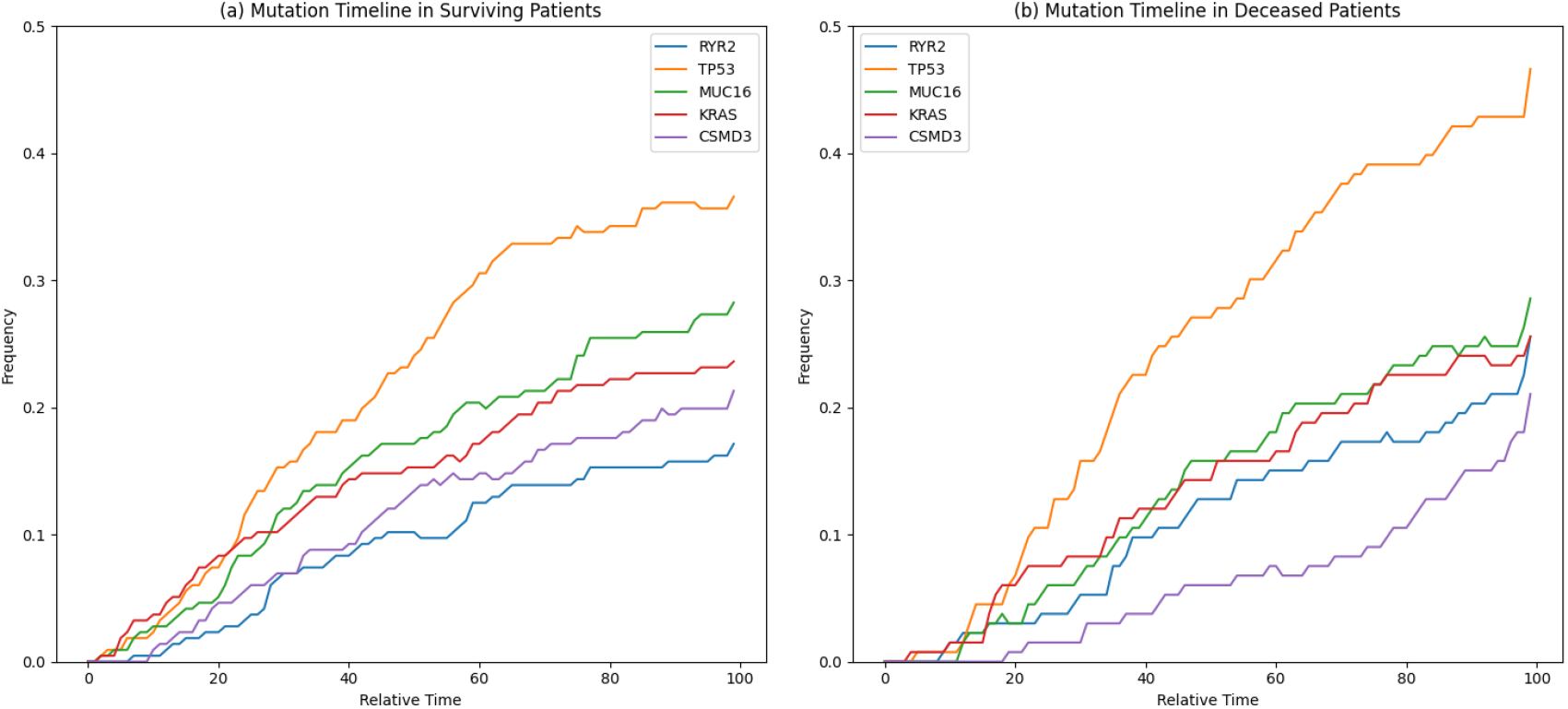
Fractions of patients with key mutations in the evolution of cancer cells for LUAD. For each subfigure, the X-axis shows the relative time of the mutation and the Y-axis is the fraction of patients with the mutation. (a) Accumulation of mutations over time in surviving patients. (b) Accumulation of mutations over time in deceased patients.

**Fig. 4:**
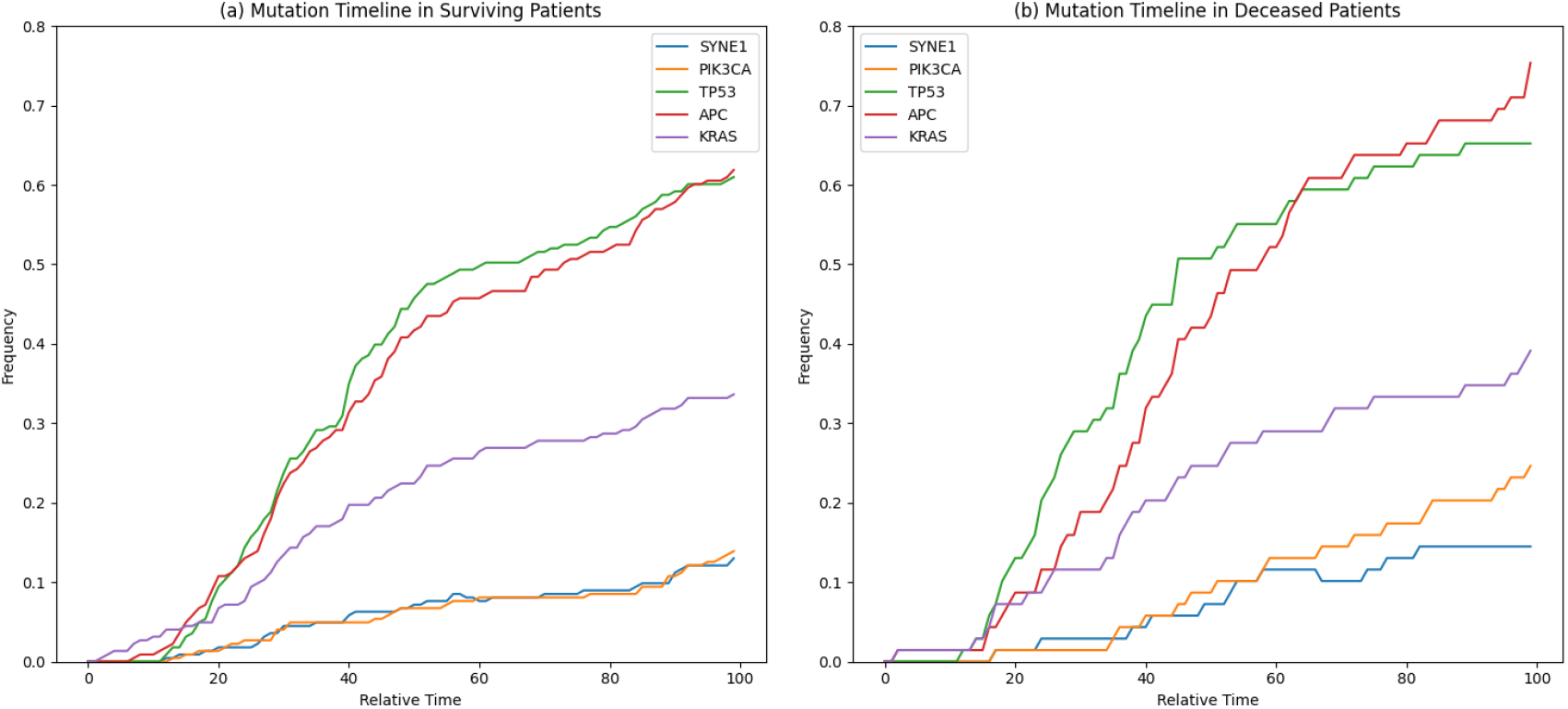
Fractions of patients with key mutations in the evolution of cancer cells for COAD. For each subfigure, the X-axis shows the relative time of the mutation and the Y-axis is the fraction of patients with the mutation. (a) Accumulation of mutations over time in surviving patients. (b) Accumulation of mutations over time in deceased patients.

To better understand how these changes at the gene level manifest as changes in predicted risk, we analyze aggregate measures of risk accumulation in Fig 5 (LUAD) Fig 6 (COAD). Fig 5(a,b) show that for LUAD there are small but not statistically significant differences between good and bad outcome groups from the root node of the tree but that these rise to significance for the most high-risk clones. Fig 6(a,b) show a qualitatively similar trend for COAD but with less pronounced differences by outcome. Fig 5(c) and Fig 6(c) show how survival time and max score relate for the two cohorts, revealing high variance patient-to-patient relative to the main separation by outcome in either cohort. Fig 5(d) and Fig 6(d) compare scores of root node versus most advanced node for each subject to give a measure of how much risk increases over the course of a cancer’s history. A more pronounced change is evident for LUAD than COAD on average, although again with high variability subject-to-subject.

**Fig. 5:**
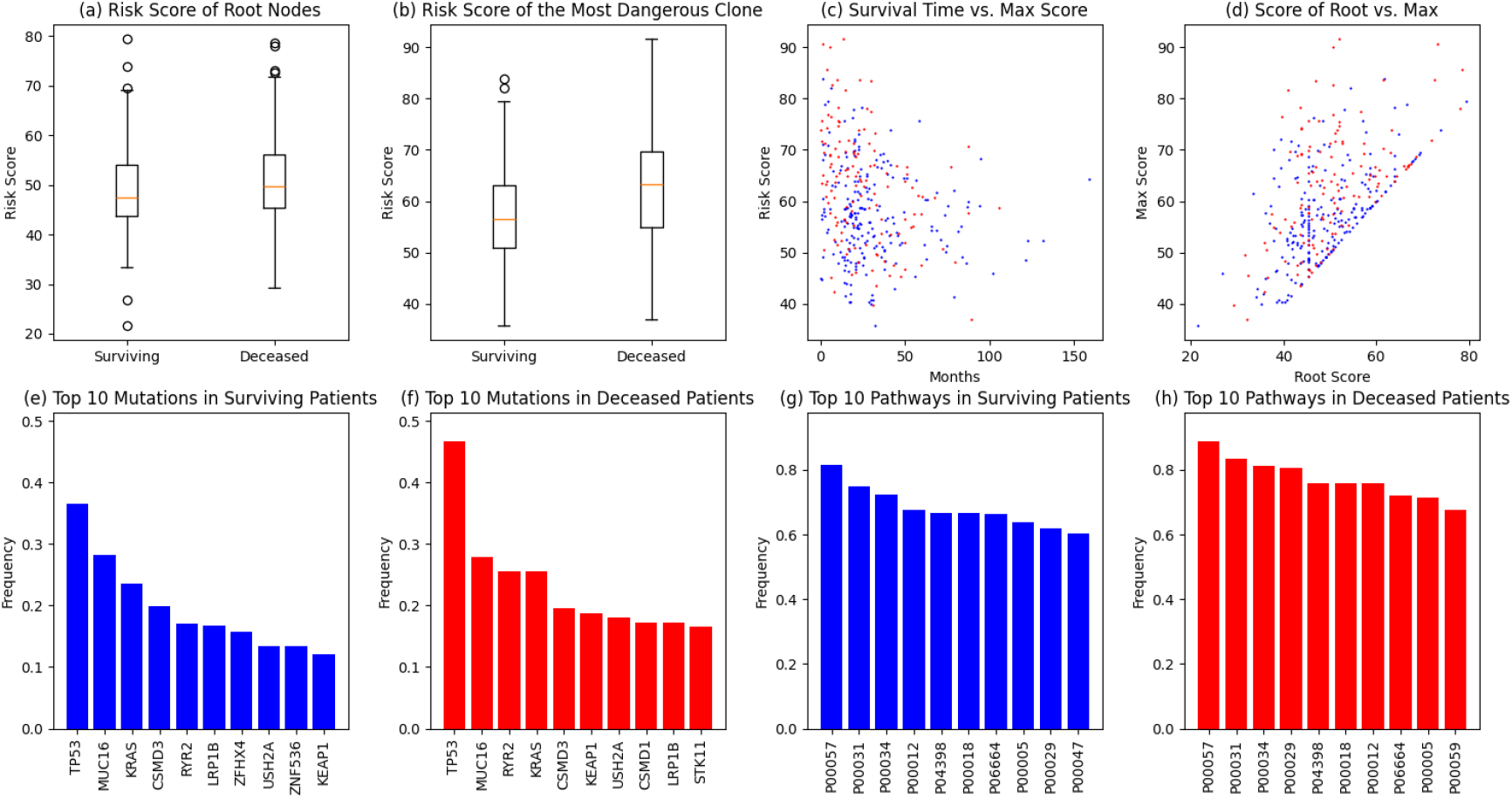
Drivers of risk evolution for LUAD.

**Fig. 6:**
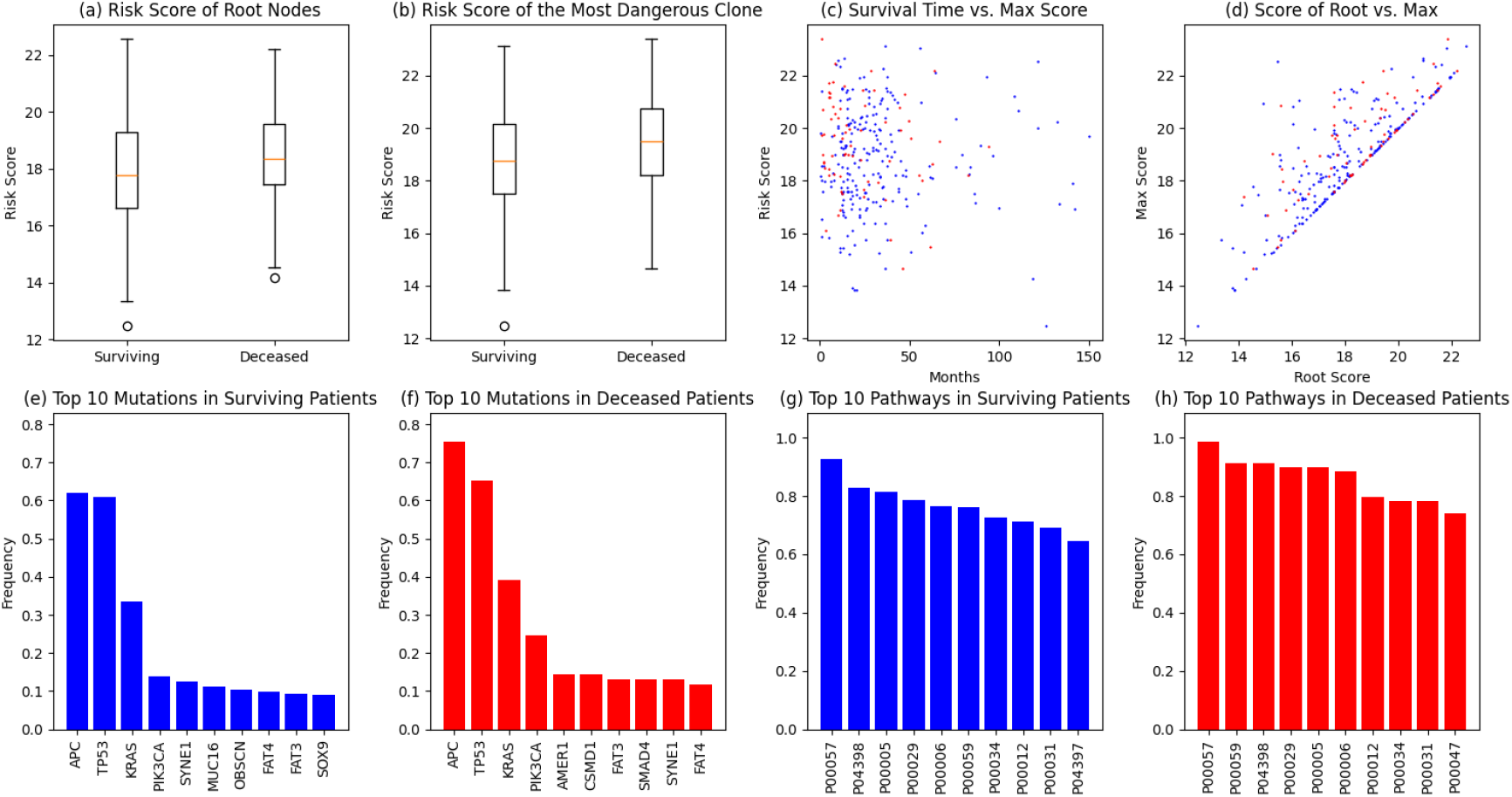
Drivers of risk evolution for COAD.

Fig 5(e-h) and Fig 6(e-h) examine overall accumulation of mutations for the top genes and pathways for each cancer, broken down by good and bad outcome cancers. Table 1 provides descriptors for the pathways identified by this analysis. In both cases, the top ranked genes and pathways are similar as are frequencies between cohorts. The high-ranking pathways themselves are largely consistent with prior expectations, for example through prominent representation of Wnt signalling, TP53 signalling, and inflammatory pathways. The most striking difference for LUAD is the relatively higher rate of TP53 mutations and somewhat high rates of mutations across the top scoring pathways in subjects with poor outcomes. COAD shows similar results with notably higher frequencies of mutations in TP53 and APC in deceased versus surviving groups and a qualitatively similar difference in pathways affected despite somewhat different pathways showing up as relevant.

**Table 1.** List of most frequently mutated pathway names to pathway IDs in both cohorts.

| Pathway ID | Pathway Name |
| --- | --- |
| P00057 | Wnt signaling pathway |
| P04398 | P53 pathway feedback loops 2 |
| P00031 | Inflammation mediated by chemokine and cytokine signaling pathway |
| P00034 | Integrin signalling pathway |
| P00012 | Cadherin signaling pathway |
| P00018 | EGF receptor signaling pathway |
| P06664 | Gonadotropin releasing hormone receptor pathway |
| P00005 | Angiogenesis |
| P00029 | Huntington disease |
| P00047 | PDGF signaling pathway |
| P00059 | p53 pathway |
| P00006 | Apoptosis signaling pathway |
| P04397 | p53 pathway by glucose deprivation |

The individual genes and pathways arising in these analyses are consistent with expectations from prior literature and the known biology of the diseases. For example, EGFR is known to be important in early lung cancers (Zhang et al., 2019) and although it does not show up as one of the five most frequently mutated genes, we can examine it individually and observe that it indeed predicted by our analysis to accumulate predominantly early in poor-outcome LUAD cancers (Figure 7). We note that some other key mutations, such as ERBB2, occur mostly through copy number alterations (CNAs) and would not be seen in this study. Other key mutations in defining cancer risk, such as TP53 (Zhang et al., 2019) have been suggested to occur more likely in the later stages of cancer evolution, although our results suggest high variability across patient populations. In contrast, key risk-driving mutations in COAD (Wu et al., 2020) appear less likely to be seen in the root node of the phylogenetic tree but rather appear to accumulate more gradually.

**Fig. 7:**
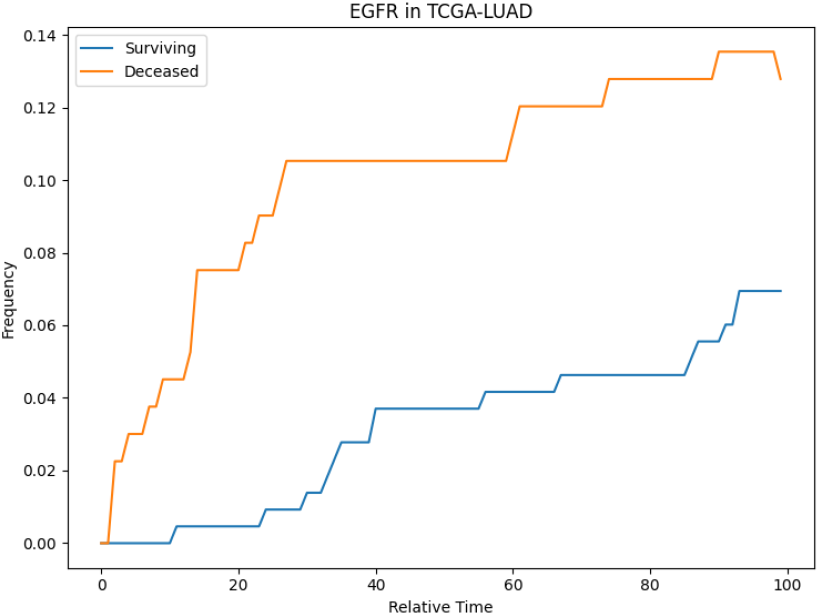
Fraction of mutated EGFR versus estimated time for LUAD.

## 4. Conclusion and Discussion

We sought to characterize how risk of progression to aggressive cancer develops over a cancer’s history. We established a model for how to answer this question from existing cancer genomic and outcome data through computational inference. We applied an implementation of that model to two cancer cohorts, revealing qualitatively similar portraits of risk accumulation although with some variability by tumor type and large variability by subject within each tumor type. The results suggest that there are average differences predictive of outcome, captured in machine learning-derived risk scores, that are present from the earliest stages of tumor development. However, these differences are dominated by high variability within each cohort. Statistically significant separation of of good and bad outcome subjects by risk score becomes possible only relatively late, an outcome that in part reflects the high variability by subject. However, this separation might be pushed back to earlier times with larger cohorts, more comprehensive variant data, or more accurate phylogeny inferences, among other factors. The results suggest that there is potential for at least subsets of patients for earlier diagnostics or interventions before cancer is commonly detected, and potentially even before it is cancer, but further advances in methodology will be needed to judge the absolute limits.

As noted in the Introduction, our goal is not to definitively answer the question the study poses about the evolution of cancer risk but rather to demonstrate that it is possible and worthwhile to answer it, in the hope of inspiring future work on the question. The biggest obstacle to answering this question better is data.

The present work used TCGA data because the nature of the question requires relatively large cohorts with consistent data types across them, and TCGA was a landmark effort that for the first time made this kind of data available. The TCGA sequencing was not designed with tumor phylogenetics in mind, though, and did not have access to many technologies available to us today. We further expect that we are missing a substantial portion of risk-driving mutations because we focus here only on SNV mutations, ignoring copy number alteration (CNA) and various kinds of structural variation (SV) mutations important in driving cancers. It is possible to map SNV, CNA, and SV mutations to a tumor lineage tree tree (Eaton et al., 2018; Fu et al., 2022), but tools for this task perform poorly on single-sample data such as is available with TCGA. Sufficiently large cohorts with multiple samples per patient, single-cell genomic data, longitudinal data, and various advances developed for studying somatic variation in non-cancerous tissues (Martincorena and Campbell, 2015) are all directions feasible now that might improve the effectiveness of future studies along these lines. How best to choose among many possible options in design of such a study is itself an emerging question for which tools are beginning to become available (Srivatsa and Schwartz, 2024). It would likewise be of interest to examine these questions in more tumor types, which would require more large cohorts than were available in the TCGA data. The computational tools used in this study are also older, in part because they were selected to be suitable for the kinds and scopes of data available and better methods for the phylogenetics and for the risk prediction aspects of this study might also be chosen or developed to accommodate richer data sources.

Finally, we consider what one would do with answers from a study such as is proposed here. Better models of cancer risk progression could have particular value to early cancer screening and public intervention efforts. As we now appreciate that somatic variation is ubiquitous (Acha-Sagredo et al., 2022) and usually asymptomatic, it is important to develop better ways to identify when seemingly harmless variation poses a risk and better respond to problems of both overtreatment and undertreatment (Kocik, 2020). Studies such as that prototyped here may also inspire better ways to respond to cancers prophylactically before they become threatening. Better characterizing the features defining a high-risk lesion may also help during cancer treatment in tailoring personalized therapies to high-risk clones within a tumor or predicting when and how a tumor under treatment might recur or become more aggressive.

## 6. Author contributions statement

This study was conceived by RS and the methodology developed by RS and KC. KC developed the code used in the study, retrieved data, and carried out the analyses. KC and RS both contributed to data interpretation and to writing of the manuscript.

## 7. Acknowledgments

This work was supported by the National Human Genome Research Institute of the National Institutes of Health under award number R01HG010589. The content is solely the responsibility of the authors and does not necessarily represent the official views of the National Institutes of Health. R.S. was supported in part by funds from Highmark Health and UPMC Enterprises through the Center for Machine Learning and Health.

